# Exonic Disruption Facilitates Antiviral CRISPR-Cas9 Activity for Multistrain HIV-1 Elimination

**DOI:** 10.1101/2021.01.14.426544

**Authors:** Jonathan Herskovitz, Mahmudul Hasan, Milankumar Patel, Wilson R. Blomberg, Jacob D. Cohen, Jatin Machhi, Daniel Stein, Evan A. Schroder, JoEllyn McMillan, Channabasavaiah B. Gurumurthy, Bhavesh D. Kevadiya, Howard E. Gendelman

## Abstract

A barrier to HIV-1 cure rests in the persistence of proviral DNA in infected CD4+ leukocytes. The high mutation rate of HIV-1 gives rise to numerous circulating strains with increased capacity for immune evasion and antiretroviral drug resistance. To facilitate viral elimination while accounting for this diversity, we propose genetic inactivation of proviral DNA with CRISPR-spCas9. We designed a library of “mosaic gRNAs” against a HIV-1 consensus sequence constructed from 4004 clinical strains, targeting the viral transcriptional regulator tat. Testing in 7 HIV-1 transmitted founder strains led, on average, to viral reductions of 82% with tandem TatD and TatE (TatDE) treatment. No off-target cleavages were recorded. Lentiviral transduction of TatDE attenuated latency reversal by 94% in HIV-infected, transcriptionally silent ACH2 T cells. In all, TatDE guide RNAs successfully disrupted 5 separate HIV-1 exons (*tat*_1-2_/*rev*_1-2_/*gp41*) providing a pathway for CRISPR-directed HIV-1 cure therapies.

**Significance Statement:** Over 38 million individuals worldwide are infected with HIV-1, which necessitates lifelong dependence on antiretroviral therapy (ART) to prevent viral replication that leads to AIDS. Efforts to rid hosts of HIV-1 are limited by the virus’ abilities to integrate proviral DNA in nuclei, mutate their genomes, and lay dormant for decades during ART treatment. We developed mosaic guide RNAs, TatD and TatE, for CRISPR-Cas9 that recognize the majority of known HIV-1 strains and inactivate 94% of proviral DNA in latently infected cells. Tandem TatDE-CRISPR inactivation of 5 viral exons (t*at*_1-2_, *rev*_1-2_, and *gp41*), which blocked HIV-1 replication for 28 days in CD4+ T cells without unwanted editing to the host genome, may serve as a viable strategy for HIV cure.

## Introduction

Defining the scientific pathways required to eliminate the human immunodeficiency virus type one (HIV-1) from its infected human host remains a global health concern for 38 million infected people. To avoid HIV-1 reactivation from latent integrated proviral DNA in CD4+ T cells and mononuclear phagocytes, lifelong antiretroviral therapy (ART) is required. ART protects against virus-induced reductions in numbers and function of CD4+ T lymphocytes and progression to the acquired immunodeficiency syndrome (AIDS) (1). Thus, a functional cure for HIV-1 is needed in order to eliminate dependence on lifelong ART.

While curative measures for HIV-1 are under development, gene editing stands alone in having the ability to remove virus independently from host antiviral adaptive immunity. Although concurrent viral elimination strategies that include broadly neutralizing antibodies (bnAbs) (2), immune modulation (3), and chimeric antigen receptor (CAR)-T cells (4, 5) show promise in mice, selection for bnAb resistant strains (6), viral rebound (3, 7), and adverse side effects (8) have been reported in monkeys and humans. Treatments notably depend on host FcR-mediated cellular killing or antigen presentation in MHC-I and, as such, rely on host immunity for viral clearance. Thus, gene therapy offers the ability block propagation of HIV-1 infection by directly targeting either host cell receptors or HIV-1 proviral DNA itself. The viral co-receptor CCR5 can be ablated *ex vivo* (9, 10) while latent viral DNA can be excised in live animals (11). Based on these results, gene therapy for HIV-1 elimination is under active clinical investigation (e.g. NCT02140944, NCT03164135, NCT03666871).

Nonetheless, applying gene therapy towards HIV-1 cure strategies has met several obstacles. One of the most significant rests in viral DNA sequence diversity. With a mutation rate of 1 in 1700 nucleotides (12), HIV-1 quasispecies easily accumulate during the course of viral infection (13). Divergent strains with increased replicative fitness evade host immunity in individuals and circulate through populations (13) as evidenced by the emergence of 11 global HIV-1 subtypes (also referred to as clades; A-L) (14). HIV-1 subtypes B and C are the most prevalent clades in Europe and the Americas (10.2%) (15) and worldwide (46%) (16), respectively. Thus, gene therapies designed to broadly target then eliminate HIV-1 must overcome strain diversity.

We posit that **c**lustered **r**egularly **i**nterspaced **s**hort **p**alindromic **r**epeat (CRISPR) approaches can meet the challenge of viral diversity. Potential proviral DNA targets of the 10 kilobase pair (kb) HIV-1 genome include the viral promoter long terminal repeat (LTR) (17–20) and viral transcription factor *tat* (21–23). CRISPR gene editing can cleave two alternatively spliced HIV-1 *tat* exons. Thus, CRISPR can simultaneously cleave *rev* and *env* that are transcribed from the same portion of the genome but in alternate open reading frames (21).

Defining the optimal gRNAs, alone or in combination, to eliminate HIV-1 infection and implementing the approach towards human trials is a challenge. *First*, computational assessments suggest that proviral targeting choices require affirmation in patient derived HIV-1 isolates. Indeed, previously published LTR-directed gRNAs likely possess limited cleavage efficiencies during natural infection (24). *Second*, gRNAs must inactivate HIV-1 transcription precluding the assembly of functional virions *en trans* from intact viral operons. *Third*, optimal combination gRNAs must minimize any emergence of CRISPR-resistant escape mutants (20, 23). While multiplexed CRISPR-Cas9 treatments were designed to account for HIV-1 strain variation (25) and minimize viral escape (21), testing against transmitted founder viral strains was not completed. *Fourth*, a CRISPR-Cas9 delivery system seeking to find and eliminate virus from its natural target cells must be employed.

Accepting each of these needs, we developed then characterized a CRISPR-Cas9 system capable of eliminating diverse proviral HIV-1 strains. The approach taken mirrors the “mosaic” HIV-1 vaccine design currently in late phase clinical trials (NCT03964415, NCT03060629) serving to immunize those persons with a high risk of acquiring infection with HIV-1 gag/pol/env antigens encoded from global composite of viral strains (26). The data offered supports the concept that CRISPR-Cas9 gRNAs designed against a multiclade HIV-1 consensus sequence, hereafter termed “mosaic gRNAs”, can prove effective. The mosaic gRNAs targeting genetically conserved and overlapping *tat/rev* exons maximized CRISPR-based attenuation of HIV-1 infection. A lentiviral transduced CRISPR-Cas9 system proved successful in its ability to block HIV-1 reactivation from latently infected CD4+ T cells. The data, taken together, brings the approach one step closer towards defining a workable strategy for HIV-1 cure.

## Results

### *In silico* design of HIV-1 *tat* mosaic gRNAs

Targeted HIV-1 gene therapy requires specific recognition of diverse proviral sequences. HIV-1 sequence diversity occurs both at the population level and during disease progression, thus lessening the effectiveness of host immune responses and promoting the emergence of antiretroviral drug resistance (27). While the 5′ LTR is a promoter for viral transcription and is susceptible to CRISPR-based HIV-1 inactivation, we reasoned that targeting the driver of LTR-directed full-length transcription, HIV-1 *tat,* would improve CRISPR therapeutic efficacy. To test this notion, a multiple sequence alignment heat map, constructed from 4004 HIV-1 strains, was utilized to observe the degree of nucleotide heterogeneity across the viral genome (**Figure 1**). Conserved regions of the HIV-1 genome include portions of flanking LTRs, gp120-encoding segments of *env*, and multiple exon overlaps (**Figure 1A**). Low entropy in HIV-1 genes encoding p6/protease, tat/rev, vpu/gp120, and nef/3’ LTR highlight sequence preservation in loci that govern multiple phases of the viral lifecycle.

**Figure 1.**
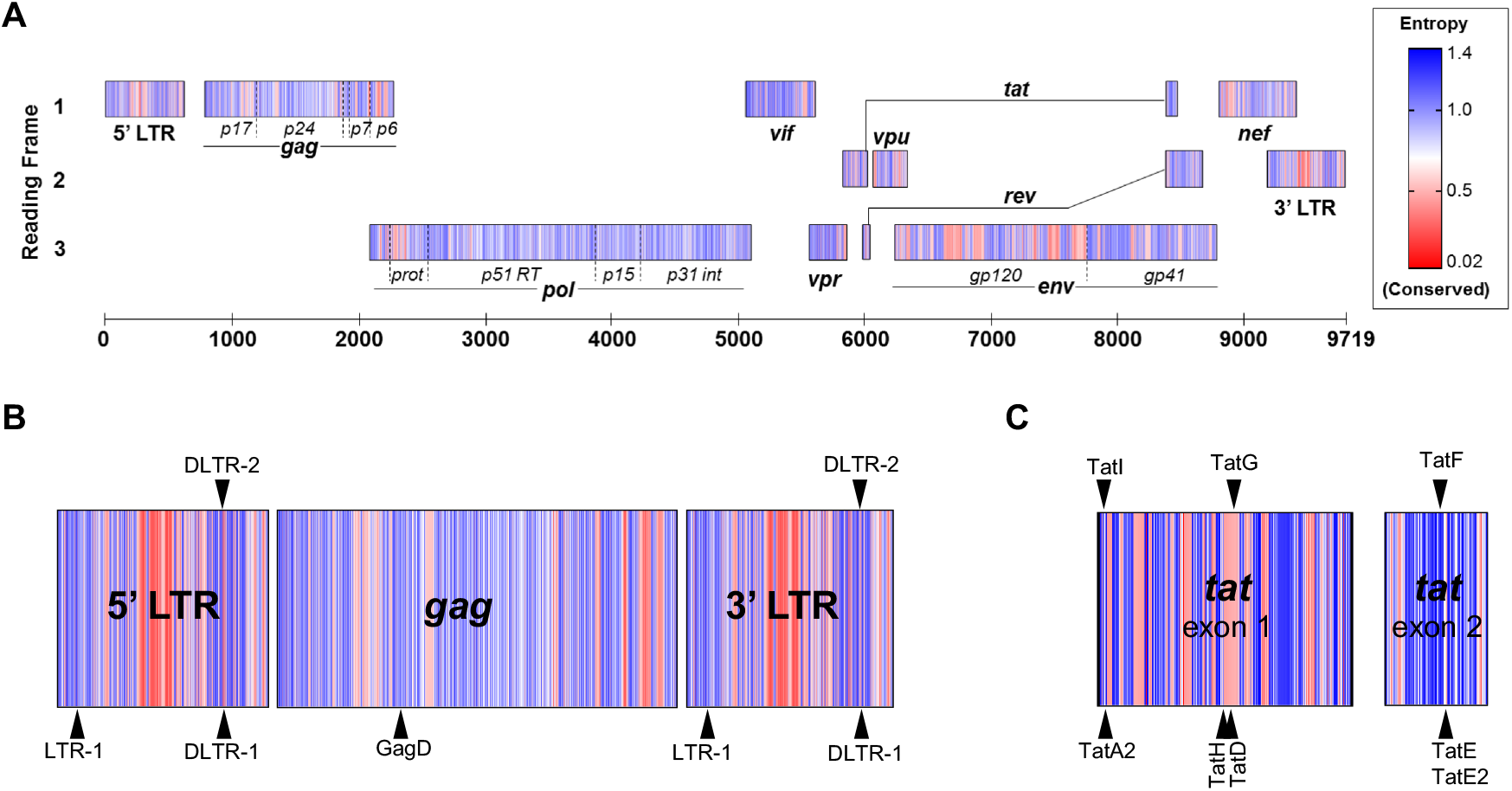
HIV CRISPR-Cas9 Mosaic gRNA Design. (*A*) Nucleotide heterogeneity of 4004 annotated HIV-1 strains is depicted in heat-map form demonstrating entropic (blue) or conserved (red) loci in all 3 reading frames. Previously reported gRNAs against LTR- and gag regions, used herein as reference controls (*B*), or novel gRNAs targeting mosaic tat sequences of HIV-1 (*C*) were designed. gRNAs against antisense or sense sequences are shown as downward or upward facing arrows, respectively.

As comparators, two CRISPR-Cas9 reference systems were adopted. *First*, LTR-1 and GagD are gRNAs that target the U3 portion of LTR and *MA* region of *gag*, respectively (**Figures 1B, S1**). LTR-1/GagD was selected as a reference control because of its utility in sterilizing one-third of HIV-1 infected humanized mice, thereby establishing proof-of-concept for a CRISPR-based HIV cure in an animal model (11). *Second*, DLTR-1 and DLTR-2 are top-rated gRNAs from a quadruplex panel designed to cleave most known HIV-1 strains with minimal off-target human genome recognition (25). Delivery of DLTR-1 and DLTR-2 gRNAs to cells remains a challenge, however, because their targeting of HIV-1 LTR (**Figure S1**) simultaneously also thwarts the synthesis of all CRISPR-transducing lentiviruses. Control gRNAs display complete 20 base pair (bp) recognition of 11-30% of curated HIV-1 strains (**Table S1**). A minimum off-target threshold of 2.266 was set for *tat* gRNA design in accordance with a CRISPR-specificity algorithm score for the DLTR-2 control.

A library of eight HIV-1 *tat*-targeting gRNAs designed for deployment with *Streptococcus pyogenes* Cas9 (spCas9) endonuclease were created. These “mosaic” gRNAs were constructed against the *tat* consensus sequence synthesized from 4004 HIV-1 strains (**Figure S1**). Five of the eight produced gRNAs target the sense strand (**Figure 1C**). Because portions of *tat* overlap with *rev* and the gp41 portion of *env*, up to three exons could be simultaneously disrupted with these gRNAs (**Figure S2**). Notably, *tat*-directed gRNAs retain full 20 bp complementarity for 6-67% of all known HIV-1 strains (**Table 1**). Duplexed mosaic gRNA treatments were found to target at least 56-62% of strains, assuming no nucleotide mismatch tolerance (**Table S2**). Taken together, these data demonstrate that gRNAs can be produced with broad HIV-1 excision potential while, at the same time, limiting off-target effects.

### *Tat/rev*-directed gRNAs suppress HIV-1 replication

To test the hypothesis that conserved gRNAs would most effectively limit viral replication, a plasmid co-transfection screen was implemented. Human embryonic kidney (HEK293FT) cells were transfected with plasmids encoding HIV-1 in the absence (untreated control) or presence of spCas9-gRNA constructs. Seven HIV-1 molecular clones, one laboratory (NL4-3) plus six clade B founder strains, were included to reflect predominant North American and European viral subtype sequence heterogeneity. Four independent co-transfection screens were performed to ensure experimental accuracy. Supernatants were measured after 72 hours for HIV reverse transcriptase (RT) activity.

The single mosaic gRNA CRISPR TatE and TatD constructs demonstrated the greatest suppression of HIV-1 replication (**Figure 2A**). TatE gRNA reduced RT activity by an average of 76% (± 6.5%, standard error of the mean (SEM)), with more than 75% reduction in 6 of 7 tested HIV-1 strains. Notably, TatE outperformed all single gRNA controls, including DLTR-2. Pearson correlation analysis evaluated whether CRISPR gRNA activity was associated with target sequence conservation (**Figure 2B**). While positive trends were observed, significance was achieved when TatE was excluded from analysis. The data support the idea that although gRNA targeting conserved genomic regions of HIV-1 proviral DNA are generally more efficacious, the TatE locus is critical to viral replication.

**Figure 2.**
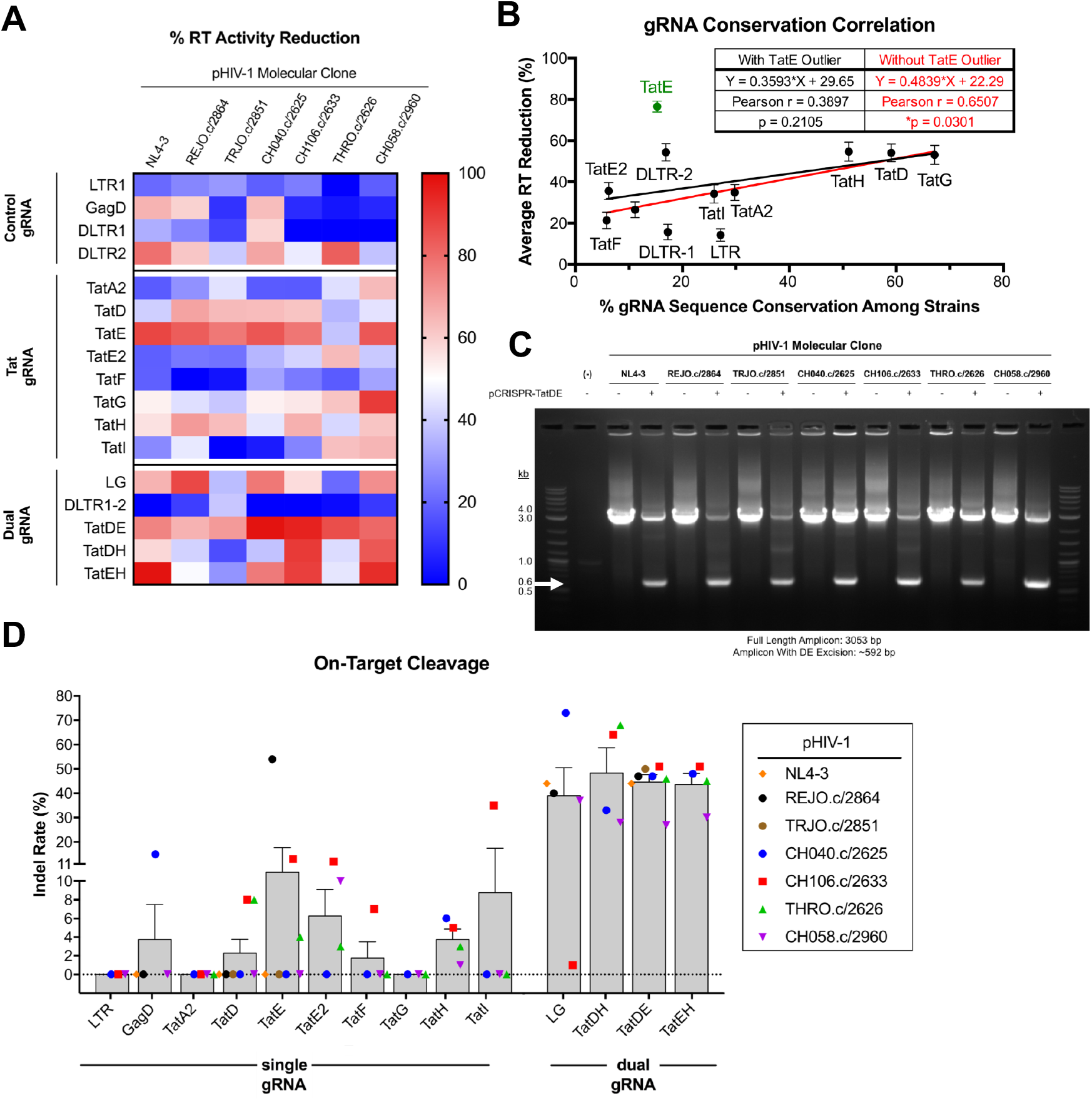
TatDE Dual gRNAs Facilitate Multistrain HIV-1 Excision. (*A*) A gRNA library was screened against multiple HIV-1 molecular clones via co-transfection of HEK293FT cells for reduction in viral replication as measured by reverse-transcriptase (RT) activity assay. (*B*) Pearson correlation between gRNA target conservation among 4004 proviral DNA sequences and average RT activity knockdown was assessed. (*C*) PCR was performed on DNA extracted from PCR amplified untreated- or CRISPR-TatDE plasmid-treated cells. White arrow indicates expected molecular size of TatDE excision bands. (*D*) PCR reaction contents were Sanger sequenced and subjected to Inference of CRISPR Edits v2.0 (ICE, Synthego 2020) to visualize nucleotide editing in the PAM/protospacer region. Data in (*A-B*, *D*) depict mean ± SEM from four independent experiments each containing biological triplicates.

We next sought to determine whether proviral DNA excision induced by two CRISPR-Cas9 gRNAs would further suppress HIV-1 replication. The noted efficacy of TatE, which targets a region of *tat* that overlaps with both *rev* and *env*, prompted us to posit that simultaneous disruptions of multiple viral exons would most drastically blunt HIV-1’s lifecycle. Plasmids were subcloned to express various combinations of the top three performing *tat*-directed gRNAs. **Table S2** summarizes the percent conservation, excision fragment length, and number of deactivated exons elicited by each pairing. Increased antiretroviral activity was observed when TatD and TatE gRNAs were co-expressed (TatDE). Viral replication in TatDE-treated cells were lowered on average by 82% (± 4.6% SEM), with a range of 65-98% across all tested viral strains (**Figure 2A**). TatDE demonstrated greater levels of viral suppression compared to LTR-1/GagD (“LG”) and DLTR-1/DLTR-2 (“DLTR1-2”) duplexed reference CRISPR controls by measured RT activity. The ~2.5 kilobase pair (kb) dropout of intervening proviral DNA was observed against all seven HIV-1 molecular clones (**Figures 2C, S3**). To confirm the accuracy of TatDE CRISPR, excision bands were Sanger sequenced and analyzed by Inference of CRISPR Edits (ICE) algorithm (**Figure S4**). Sequencing data qualitatively depict nucleotide degeneracy in both target loci, with insertion/deletion (indel) mutations at a rate of 25-51% in excision bands. Pairing our two most conserved efficacious gRNAs, TatDH, and of our largest excising duplex, TatEH, proved inferior to 5-exon inactivating TatDE in suppressing RT activity. Taken together, these data support the hypothesis that disrupting a maximal number of viral exons by CRISPR-Cas9 leads to the greatest suppression of HIV-1 replication.

### On-target CRISPR-Cas9 TatDE specificity

We next evaluated the specificity of our *tat*-directed CRISPR-Cas9 by comparing the degree of proviral genome editing to that observed in off-target loci. ICE analysis performed on Sanger sequences from these reactions revealed that dual gRNA systems improve indel mutation rates to >40% as compared to <10% seen with single gRNA controls (**Figure 2D**). TatDE therapy resulted in an average of 45% (± 3.0% SEM) gene modification rate with 40-50% editing in 6 of 7 tested strains, supporting TatDE’s ability to deactivate a broad variety of HIV-1 proviral species.

To ensure TatDE cleavage is restricted to HIV-1 proviral DNA, we investigated the potential for aberrant indels at off-target loci in the human genome. These covered 5 possible loci recognized by each TatD, TatE, LTR-1, and GagD. At all putative off-target positions for TatD and TatE, no editing was observed (**Table S3**). This was sustained regardless of whether cells were treated with single or dual gRNAs. Similarly, off-target analysis by ICE for LTR-1 and GagD failed to demonstrate indel mutations in host genes (**Table S4**). Seven of 10 possible off-target regions for TatDE fell in intronic regions whereas only 2 of 10 for LTR-1/GagD were noncoding. These results reinforce the notion that TatDE CRISPR is not more likely to affect host gene expression than other HIV-1 CRISPR systems.

### TatDE CRISPR-Cas9 eliminates latent proviral HIV-1

Having demonstrated the ability of TatDE CRISPR to suppress plasmid-encoded viral transcription, we next investigated whether latent HIV-1 could be excised in leukocytes. For these experiments, TatD, TatE, or non-targeting control gRNAs were cloned onto a lentiviral-CRISPR expression plasmid. To first assess CRISPR-Cas9 function, ACH2 T cells and U1 promonocytes bearing 1-2 copies of latent HIV-1 proviral DNA copies per cell were transfected with cloned lentiviral-CRISPR constructs. Subsequent stimulation of ACH2 T cells with TNFα induced viral rebound in control- and LTR-1/GagD-treated cultures but not in those receiving single- or dual TatD plus TatE treatments (**Figure S5**). U1 promonocytes were equally responsive to LTR-1/GagD and TatD/TatE treatments (**Figure S6**). HIV-1 proviral DNA excision was present in both cell types and confirmed by sequencing. These data in aggregate validate the TatDE CRISPR-Cas9 system for lentiviral delivery.

We next ascertained whether lentiviral transduction of TatDE CRISPR-Cas9 could halt HIV-1 induction from latently infected ACH2 T cells. Transgene expression was measured by reverse-transcriptase quantitative PCR (RT-qPCR) and determined to be above detection limits (**Figure 3A**). Dual TatD/TatE lentiviral treatments blocked TNFα-induced stimulation. Cotransduction of TatD/TatE at MOIs of 10 and 1 significantly reduced the release of HIV-1 into culture supernatants by 80% and 94%, respectively (**Figures 3B-C**). We also evaluated cell DNA by nested PCR for TatDE HIV-1 excision (**Figures 3E, S7**). A predictable ~525 bp excision band was present for cells co-transduced at a MOI of 1, supporting the reduction in RT activity at this MOI. Shorter amplicons were observed with MOI 0.1 treatment, however the shortened amplicons disappeared in stimulated conditions. Paired with the RT activity results (**Figure 3D**), it is likely that transduction at a low MOI was subtherapeutic, as few cells bear the CRISPR-induced indel. Conversely, we cannot exclude the possibility that the efficacy observed at MOI 10 in the absence of excision bands results from cytopathicity encountered with high levels of lentiviral treatment. These results cumulatively support the utility of lentiviral delivery of TatDE CRISPR-Cas9 in eliminating latent proviral HIV-1.

**Figure 3.**
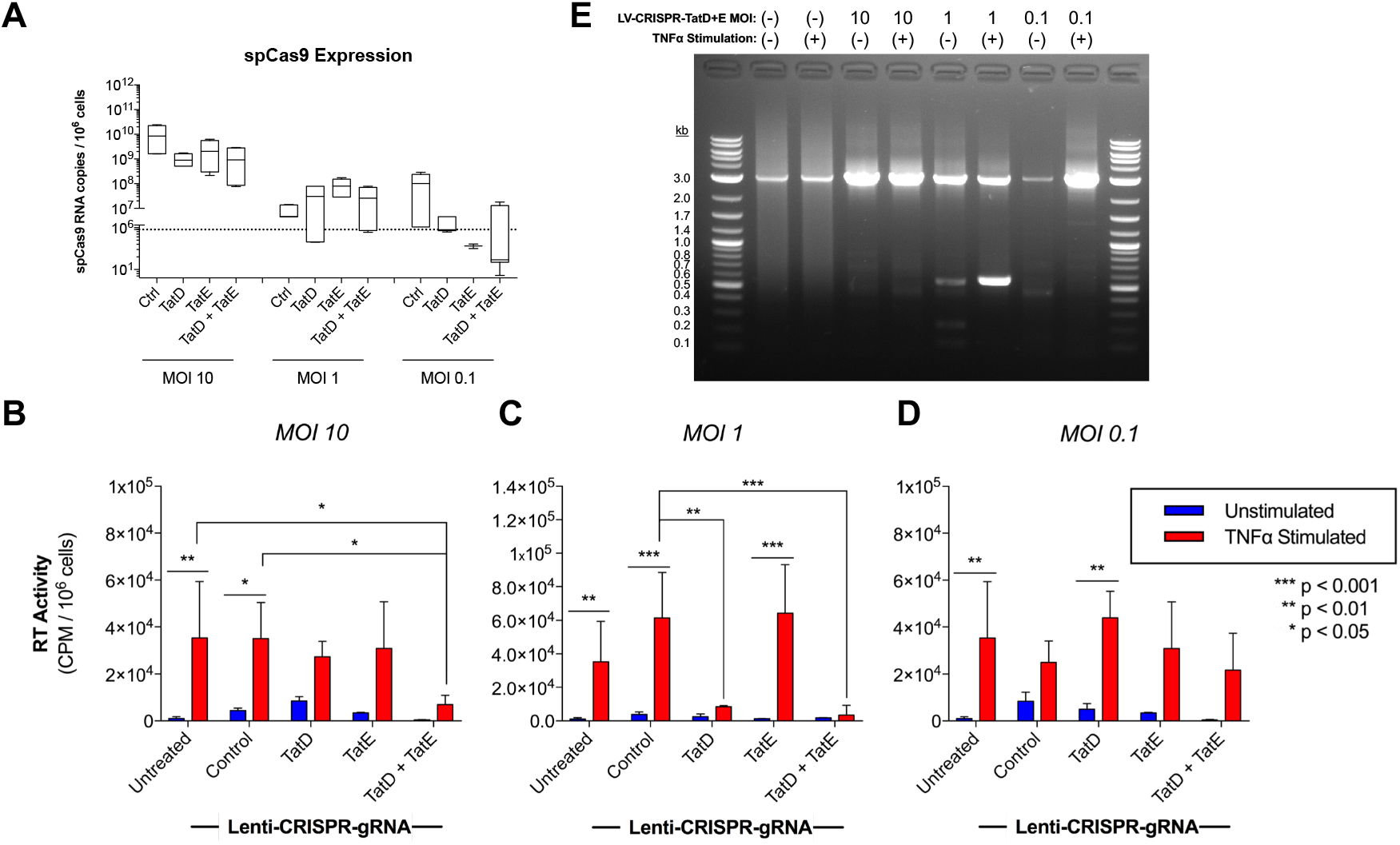
Lentiviral TatD + TatE CRISPR co-transduction Inactivates Latent HIV-1. ACH2 T cells bearing 1 copy of HIV-1 proviral DNA were transduced with lentivirus bearing *spCas9-gRNA* transgene at multiplicities of infection (MOI) 10, 1, or 0.1. After 72 hours, cells were stimulated with TNFα (15 ng/mL) for 72 additional hours. (*A*) *spCas9* expression as measured by RT-qPCR. (*B-D*) RT activity assay from culture supernatants. (*E*) Nested PCR for proviral DNA excision wherein unedited amplicons are 2986 bp and CRISPR-edited amplicons are approximately 525 bp, depending on insertion / deletion mutagenesis. White arrow indicates expected molecular size in presence of TatDE excision. Significance assessed by two-way ANOVA.

### Exonic disruption of the *tatE* locus attenuates HIV-1 replication

As a potential mechanism to explain TatE’s high antiretroviral activity despite low sequence conservation, we investigated how disruption of different numbers of viral exons affects HIV-1 replicative fitness. Three HIV-1_NL4-3-Δnef-eGFP_ non-frameshift point mutants were created to parallel CRISPR mutation profiles (**Figure 4A**), in which two-(ΔTatD), three-(ΔTatE), or five (ΔTatDE) exons were altered (**Figure 4B**). Resulting virions were imaged by transmission electron microscopy (TEM; **Figure 4C**) and titered by RT-qPCR. The size of the HIV-1 *tat* mutants ranged from 110.6-150.5 nm in diameter, closely approximating that of wildtype HIV-1 measured at 125.4 nm. Likewise, the HIV-1 *tat* mutants’ nearly spherical appearance and inner conical capsid matched the morphology of wildtype control virus. Next, CD4+ T lymphocytes were infected at a MOI of 0.1. At 10 days following HIV-1 infection, significant differences in RT activity were detected in culture supernatants (**Figure 4D**). HIV-1_NL4-3-Δ*tatD*_ displayed nearly equivalent levels of viral replication as wildtype virus, while mutants bearing ≥ 3 disrupted exons were significantly lower in RT activity compared to unmutated control. As CD4+ T cell proliferation rates differed after 10 days due to HIV-1 induced cytopathicity, the percent of cultured HIV-1 infected cells was measured by flow cytometry during a 28-day time course (**Figure 4E**). Whereas HIV-1_NL4-3-Δ*tatD*_ proliferation approximated that of control HIV-1, HIV-1_NL4-3-Δ*tatE*_ and HIV-1_NL4-3-Δ*tatDE*_ remained at or around baseline for four weeks. These data suggest that the locus targeted by TatE gRNA is critical for maximal CRISPR activity. In aggregate, we conclude that the high efficacy of TatDE CRISPR-Cas9 therapy against numerous HIV-1 strains results from disrupting five viral exons simultaneously.

**Figure 4.**
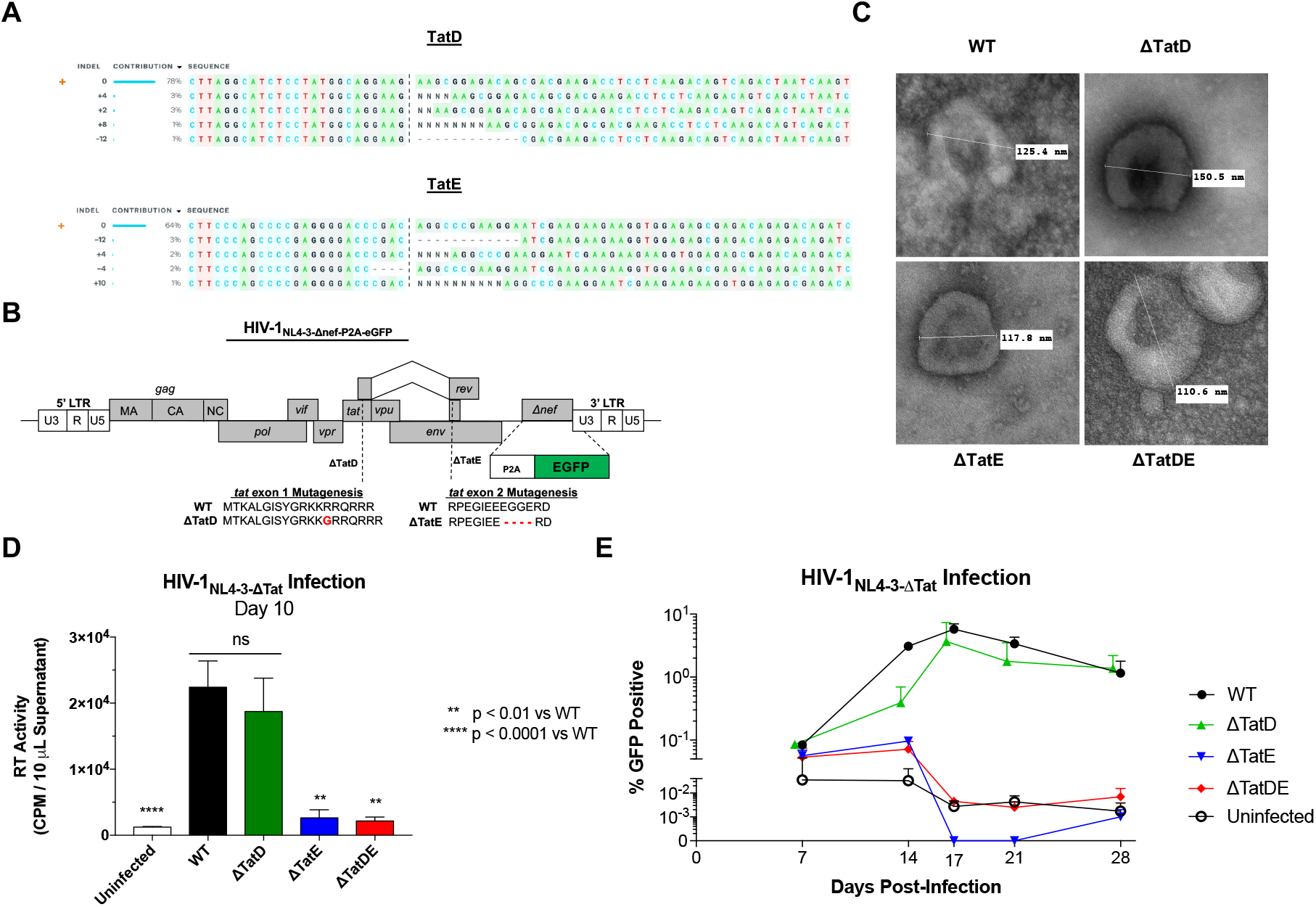
Exonic Disruption Compromises HIV-1 Replicative Fitness. (*A-B*) Insertion / deletion profiles among the most-efficacious single gRNAs from a co-transfection screen were assessed by Synthego ICE v2.0 algorithm. Highest frequency insertions or deletions were selected for subsequent non-frameshift site-directed mutagenesis of HIV-1_NL4-3-Δnef-eGFP_ encoding plasmid. (*C*) Transmission electron micrographs of single- or dual-*tat* mutants. Spherical diameter measurements were taken at the time of imaging (inset). (*D-E*) CEMss CD4+ T cell lines were challenged at MOI 0.1 with HIV-1 NL4-3-Δtat-Δnef-eGFP and assayed at designated time points for RT activity (*D*) or by flow cytometry for % GFP-positive cells (*E*).

## Discussion

Over 79,000 unique human- or simian immunodeficiency virus (HIV/SIV) sequences spanning eleven global subtypes have been identified (14). The presence of a 6% mutation rate even among elite controllers highlights the presence of a multitude of intrapatient HIV-1 quasispecies (28). These considerations led us to design mosaic CRISPR-Cas9 guide gRNAs against conserved and multi-exon regions of a consensus HIV-1 sequence to facilitate viral elimination. Herein, we introduce exonic disruption as a guiding principle for CRISPR therapeutics as a required link to control residual viral infection

Our data establish an association between *tat/rev* target DNA conservation and CRISPR-Cas9 efficacy across multiple HIV-1 transmitted founder strains (**Figure 2B**). The impact of this finding is twofold. *First*, it validates our mosaic gRNA approach as a tenable means of using CRISPR to inactivate genetically heterogenous viral targets. *Second*, the correlation reveals that conserved *tat/rev*-directed gRNAs outperform LTR-based reference controls. Given that these were selected for their potential to recognize divergent HIV-1 quasispecies in patients (25) and facilitate elimination from infected hosts (29), the comparative superiority of TatDE represents an advance in CRISPR therapy as a strategy for HIV-1 elimination.

A second advantage of the TatDE mosaic gRNA approach is that it identifies exonic disruption as a potential mechanism for antiretroviral efficacy. Assessed variables include CRISPR on-target cleavage efficiency, target DNA conservation, excision length, and numbers of disrupted exons during CRISPR-Cas9 treatment. Non-frameshift site-directed mutagenesis of 3 or more exons at the TatE locus, but not the 2 exons at TatD, significantly suppressed viral replication. In our gRNA co-transfection screen, the most broadly active CRISPR treatments included *tat/rev/gp41*-directed TatE. One implication that follows is that transmission of TatE-directed CRISPR may help overcome low potency (30) and viral escape (31, 32) otherwise associated with single gRNA treatment of HIV-1. These results also suggest that cleavage at other multiexon locations along the HIV-1 genome, such as *gag/pol* and *nef/*LTR, or *tat/rev/gp41* by other Cas nucleases (20, 33) may afford broader antiviral coverage. Thus, exonic disruption paired with target DNA conservation appear integral in generating effective HIV-1 CRISPR therapeutics.

Delivery of HIV-1 specific CRISPR payloads remains one final parameter yet to be optimized prior to human administration. Current *ex vivo* approaches ablate the HIV-1 co-receptor CCR5 with CRISPR-transducing lentiviruses (34, 35). However, the 5% editing efficiency in transduced human hematopoietic stem cells failed to render viral suppression upon antiretroviral drug withdrawal (34). These findings informed our strategy to deploy TatDE CRISPR via a highly efficient viral vector in leukocytes. With lentiviral transduction at a MOI of 1, the TatDE CRISPR transgene was detected at an average of 33.2 *spCas9* RNA copies per AHC2 T cell, with a 94% reduction in RT activity and clear viral excision (**Figure 3**). In contrast, a single broad spectrum gRNA targeting LTR transduced at this same level still allowed viral rebound in ~20% latently infected JLat cells (36). Other studies have achieved >90% reduction in HIV-1 reactivation and prevention of viral escape when CRISPR-Cas9 was lentivirally transduced in T cells at MOIs 10-15 (21, 23, 37). These differences highlight the likelihood that TatDE CRISPR-Cas9 maintains greater potency against HIV-1 compared to other CRISPR-Cas9 therapies.

A possible critique of our vector choice is that patients will likely already be receiving combination antiretroviral drugs that inhibit lentivirus’ ability to transduce CRISPR. Nonetheless, latently infected JLat cells, uninfected- and HIV-1-positive patient blood mononuclear cells have been treated successfully by lentivirus transduction in the presence of antiretroviral drugs (29, 37). This sets a stage for future optimization of dosage and ART timing to enable CRISPR delivery by lentivirus in HIV-1 infected patients. Phase I clinical trials evaluating the safety of lentiviral-transduced CRISPR-Cas9 are ongoing for B-cell acute lymphoblastic leukemia (B-ALL; NCT04557436) and HIV-1 patients in the context of hematological malignancies (NCT03164135). Thus, our ability to potently deactivate HIV-1 with lentiviral vectors may prove essential in avoiding adverse events with TatDE CRISPR therapy.

Selectivity of the CRISPR-Cas system for HIV-1 proviral DNA is a key factor in assessing our treatment’s therapeutic window. No off-target edits were observed with TatDE treatment in the 10 loci surveyed by a Sanger sequencing-based algorithm. By contrast, ICE data demonstrate a mean 45% editing efficiency of proviral DNA template with TatDE CRISPR. This led to detectable excision of ~2.5 kb of viral DNA and to nearly complete abolition of viral release. These findings are aligned with reports that ~10% and 39-63% editing efficiencies from multiplex endonuclease treatments resulted in more than 90% knockdown in herpes simplex virus and HIV-1, respectively (23, 38). Dysregulation of positive feedback loops, as in the case of *tat/rev*-directed CRISPR-Cas9, and the formation of dominant negative proteins encoded after gene editing may explain the disproportionately high efficacy in each of these cases. The HIV-1 proviral genomes isolated from elite controllers display large deletions of *env* sequences flanked by *tat/rev* amongst other regulatory genes (39). It is postulated that defective virus in elite controllers primes the host immune system to clear cells bearing replication-competent HIV-1 proviral DNA (39). Therefore, we posit that TatDE CRISPR therapy may similarly compromise latent HIV-1 sequences thereby inducing immune-mediated clearance *in vivo*.

Planned studies will serve to address future needs in translating this strategy for human use. TatDE’s multistrain potential will be further validated against additional HIV-1 strains and clades for antiviral activity. We will also interrogate the lack of off-target editing within the human genome by employing CIRCLE-seq (40, 41), Guide-seq (36, 42) and whole genome sequencing (29) to our TatDE CRISPR-treated samples. Finally, we plan to further characterize TatDE’s mechanism of action in terms of its ability to downregulate full-length viral transcripts due to *tat*, diminish *rev*-associated nuclear export, and impede gp41-mediated viral entry. These additional studies should clarify the value in expanding our conserved, multi-exon targeting CRISPR library to other Cas endonucleases.

To conclude, the combination of “**d**iverse” strain-inactivating Tat**D** and “tri-**e**xon”-directed Tat**E** gRNAs prompted maximal suppression of HIV-1 replication by CRISPR-Cas9. Further refinement of these methodologies may advance the utility of gene therapy in eliminating latent viral infection from infected hosts across the globe.

## Materials and Methods

Detailed protocols for materials and methods are provided in Supplementary Information.

### Cloning and transfection

CRISPR plasmid constructs containing mosaic gRNAs were cloned by T4 ligase oligonucleotide insertion (**Table S5**) in px333 (Addgene #64073) or pLentiCRISPR-RFP657 (Addgene #75162) and transformed into STBL3 *E. coli*. In Fusion HD Cloning (Takara #638920) was utilized to synthesize *tat* mutants on pHIV-1_NL4-3-Δnef-eGFP_ background. Maxiprep purified plasmids were transfected into HEK293FT cells using PEI or into ACH2 / U1 leukocytes via TransIT-Jurkat reagent (Mirus #2120). Supernatants were collected 72 hours after media replacement (PEI) or plasmid transfection (TransIT-Jurkat).

### Virus, transduction, and viral outgrowth assays

HIV-1_NL4-3-Δnef-eGFP_ and HIV-1_NL4-3-Δnef-eGFP-Δtat_ were synthesized by reverse-genetics from supernatants of transfected HEK293FT cells. Supernatants were passed through 0.45 μm filters and ultracentrifuged to purify and concentrate virus. HIV-1 viral stocks were titered by RT-qPCR (Takara #631235) to determine viral RNA copies / mL. Multiplicities of infection (MOI) for HIV-1 infection of CEM-SS T cells (NIH ARP #776) were calculated as viral RNA copies / cell. CRISPR-transducing lentivirus was commercially prepared at the University of Iowa Viral Vector Core Facility and titered using digital droplet PCR. ACH2 T cells (NIH ARP #349) were transduced with CRISPR-encoding lentivirus at designated MOIs such that MOI 1 = 1 transducing unit per cell. ACH2 and U1 (NIH ARP #165) were stimulated 3 days post transfection or transduction with recombinant tumor necrosis factor alpha (TNFα; 15 ng / mL) or phorbol 12-myristate 13-acetate (PMA; 20 ng / mL) for 3 additional days followed by experimental harvest.

### (RT-q)PCR

DNA was harvested from experimental pooled biological triplicates using NucleoSpin Tissue XS Micro kit (Machery-Nagel #740901). PCR for proviral DNA excision was run in 25 μL reactions using PrimeTime Gene Expression Master Mix (IDT #1055772) using 150 ng template DNA and 35 cycles or 15 cycles plus 35 cycles for nested PCR. Off-target PCRs were performed with 150 ng template DNA in 25 μL reactions using AmpliTaq Gold 360 master mix. Primer sequences for all reactions are provided in **Table S6**. Gel extracted PCR amplicons and total PCR contents were sequenced and subjected to Synthego Inference of CRISPR Edits (ICE) Analysis v2.0 (https://ice.synthego.com/#/).

RNA was harvested from pooled biological triplicates using TRIzol™ solution (Invitrogen #15596026), DNase I digested (Zymo Research #E1010), and converted to cDNA (Thermo Scientific #AB1453B). RT-qPCR was performed in technical triplicates using standard curve absolute quantitation for *gapdh* (1 diploid genome = 6.6 picograms DNA extracted from ACH2) and *spCas9* (pLenti-CRISPR-RFP657; 10^9^ copies equal 12.01 ng according to MW 7.22 × 10^6^ Da). Reactions were performed in 10 μL volumes containing 1 μL cDNA templated, 5 μL PrimeTime Gene Expression Master Mix, 500 nM of each primer (5′-CCCAAGAGGAACAGCGATAAG-3’; 5′-CCACCACCAGCACAGAATAG-3’), and 0.5 μL 20x GAPDH endogenous control probe (Applied Biosystems #4333764T) or 200 nM *spCas9* probe (56-FAM/ATCGCCAGA/ZEN-/AAGAAGGACTGGGAC/3IABkFQ).

### Reverse transcriptase activity assay

Culture supernatants were assayed for RT activity by surveying for ^3^H-thymidine incorporation (3H-TTP; Perkin Elmer #NET221A001MC). Briefly, 10 μL of culture supernatants were spiked into 96-well round bottom plates, digested for 15 minutes with 10 μL solution A (100 mM Tris-HCl pH 7.9, 300 mM KCl, 10 mM DTT, 0.1% NP-40), then reacted with 25 μL solution B (50 mM Tris-HCl pH 7.9, 150 mM KCl, 5 mM DTT, 15 mM MgCl2, 0.05% NP-40, 0.250 U/mL oligo dt pd(T)12-18), 10 μCi / mL 3H-TTP (4 μL / mL; 3H-deoxythimidine 5′-triphosphate, tetrasodium salt, (methyl-3H)). After 20 hours incubations, 50 μL ice cold 10% tricholoroacetic acid was added to plates, vacuum filtered across 96-well MicroHarvest Plates, and red by scintillation counting using TopCount Scintillation counter (Perkin Elmer).

### Statistical Analyses

Experiments were un with biological replicates (n = 3 or 4) in at least two separate trials. Data represent mean ± standard error of the mean (SEM). Pearson correlation, one-way ANOVA with Dunnet correction, and two-way ANOVA with Sidak correction for multiple comparisons were calculated using GraphPad Prism v7.0a for Mac OS X (GraphPad Software, San Diego, California USA, www.graphpad.com).

## Acknowledgments

This work was supported in part by the National Institutes of Health Grants P01 DA028555, P30 MH062261, R01 MH115860, R01 NS034249, R01 NS036126, R01 MH121402, T32 NS105594 (to H.E.G), UNMC Assistantship / Fellowship Program (to J.H. and M.H.), Nicholas Badami Fellowship (to J.H), and the Carol Swartz Emerging Neuroscience Fund (to H.E.G). The authors thank Drs. Beat Bornhauser (U. of Zurich), Kamel Khalili (Temple University), and Won-Bin Young (U. of Pittsburgh) for kindly providing plasmids used in this project. Gratitude is also extended to Nicholas Conoan and the UNMC Electron Microscopy Core for their assistance. A final word of appreciation is directed to Daniel Chadash and the engineering team at Genome Compiler for making open access cloning software freely available online.

