## Supplemental Information for "Exonic Disruption Facilitates Antiviral CRISPR-Cas9 Activity for Multistrain HIV-1 Elimination"

**Supplementary Information Text**

**Supplementary Methods.**

***Escherichia coli* (STBL3, Stellar)**

*E. coli* (STBL3, Stellar) cells used for transformation of stock or cloned plasmids were streak plated on selective antibiotic-containing (ampicillin, 100 µg/mL OR kanamycin, 50 µg/mL) Luria Bertani (LB)-agar plates and incubated 16 hours at 37°C. Single clones were then inoculated in LB broth containing appropriate selective antibiotic and cultured for 16 hours at 32°C with orbital shaking at 185 revolutions per minute (rpm). Selected clones were stored in 50% glycerol/water at -80°C.

**Human cell culture**

HEK293FT cells were grown in DMEM containing 10% v/v fetal bovine serum (FBS), 100 units / mL penicillin and streptomycin (PenStrep), 0.5 mg / mL Geneticin®, 1% v/v MEM non-essential amino acids (MEM NEAA), and 1 mM sodium pyruvate. Adherent HEK293FT were detached from culture plates using 3 mL Trypsin-EDTA (0.25%) and passaged at 3 x $10^6$ cells in 15 mL growth medium every 3-4 days. CEMss CD4+ T cells, HIV-1 latently infected ACH2 T cells, and HIV-1 latently infected U1 promonocytes were cultured in RPMI containing 10% v/v FBS, 100 units / mL PenStrep, and L-glutamine. Suspension CEMss, ACH2, and U1 cells were maintained at 0.2 - 1 x $10^6$ cells / mL by passaging in 15 mL growth medium every 3-4 days. Cells were propagated in vented cap 75 $cm^2$ U-shaped flasks at 37°C / 5% $CO_2$.

***In silico* HIV-1 Sequence Conservation and guide RNA (gRNA) Design**

To visualize conservation of nucleotides through the HIV-1 genome, a DNA multiple sequence alignment was completed for each LTR or exon according to the HXB2 reference strain (https://www.hiv.lanl.gov/content/sequence/HIV/MAP/landmark.html) from the Los Alamos National Library (LANL) HIV sequence database (https://www.hiv.lanl.gov/) using all complete sequences available through 2018. The exported FASTA files were input to WebLogo 3 (http://weblogo.threeplusone.com/create.cgi) and the resulting plain text data table of positional entropy was graphed in heat-map form. Consensus sequences lacking gaps for the entire HIV-1 genome and *tat* using the LANL multiple sequence alignments were generated using SnapGene® software (GSL Biotech). CRISPR gRNAs for HIV-1 *tat* were designed using CHOPCHOP v3 (1) (https://chopchop.cbu.uib.no) and Broad Institute GPP sgRNA Designer (https://portals.broadinstitute.org/gpp/public/analysis-tools/sgrna-design) using *tat* consensus sequence as the specified target and human GRCh38 as the host organism. The top 35 gRNA candidates were inspected in WebLogo to identify the 8 gRNAs with the highest degrees of conservation among HIV-1 strains. Guide RNA conservation percentages for the full 20 basepair target were also determined using SnapGene®. The specificities of *tat*- and control-targeting gRNAs were predicted using Broad Institute GPP sgRNA Designer and CRISPR-OFF webserver v1.1 (https://rth.dk/resources/crispr/crisproff/). Putative off-target loci for top gRNA candidates were selected from top-ranked candidates in CRISPR-OFF and Cas-OFFinder (http://www.rgenome.net/cas-offinder/) tools.

**Cloning**

CRISPR U6::gRNA-U6::gRNA-CBh::spCas9 "all in one" px333 (Addgene plasmid # 64073) and pLentiCRISPR-RFP657 (Addgene plasmid # 75162) were restriction enzyme digested using BsaI, BbsI, or Esp3I as appropriate. Digested vectors were extracted from 1% agarose / TAE using GeneJET gel extraction kit (Thermo Scientific). Guide RNA recognition domain inserts flanked with *cacc or caaa* sticky ends were generated by primer annealing in 10 mM Tris pH 8.0, 50 mM NaCl, 1 mM EDTA using thermal cycling touchdown (95°C – 25°C, Δ1°C, 45 seconds per stage). Vector and insert were ligated using T4 ligase reacted for 16 hours at 4°C, followed by heat inactivation at 65°C for 15 minutes. pHIV-$1_{NL4\text{-}3\text{-}\Delta tat}$ mutants were generated by site-directed mutagenesis of pHIV-$1_{NL4\text{-}3\text{-}\Delta nef\text{-}eGFP}$ (a gift from Dr. Won Bin Young, University of Pittsburgh, PA; and Dr. Kamel Khalili, Temple University, PA) using In-Fusion cloning (Takara) according to manufacturer's specifications and recommended primer pairings. Primers used for insertional and mutagenesis cloning are summarized in **Table S5**. CRISPR plasmid constructs and HIV-1 molecular clones pHIV-$1_{NL4\text{-}3\text{-}\Delta nef\text{-}eGFP}$, pCH040.c/2625, pCH106.c/2633, pTHRO.c/2626, pREJA.c/2864, pTRJO.c/2851, and pCH058.c/2960 were transformed into STBL3 *E. coli*. Mutant pHIV-1$_{NL4-3-\Delta tat}$ were transformed in Stellar Competent *E.* coli. Plasmids were extracted via miniprep and Sanger sequenced to confirm cloning success or via maxiprep for subsequent transfection.

**Transfection**

HEK293FT cells were seeded 24 hours prior to transfection in DMEM growth medium lacking geneticin at 5 x $10^5$ cells / well in 12-well culture plates (px333-pHIV-1 co-transfection screen) or 6 x $10^6$ cells / 10-cm culture dish (stock HIV-1 generation). Four hours prior to transfection, medium was changed to OptiMEM containing 250 µM chloroquine diphosphate (1 mL per well or 8 mL per 10-cm dish). Polyethyleneimine (PEI; linear MW 25,000 Da) stock (1 mg/mL) was diluted in OptiMEM to account for a 3:1 w/w PEI:DNA ratio. For gRNA co-transfection screening, 1.5 µg px333 plus 1.5 µg pHIV-1 were diluted in 150 uL OptiMEM, which was then further diluted with 150 uL OptiMEM / PEI. Viral stock transfections were similarly prepared using 10 µg pHIV-1 diluted in 1.5 mL OptiMEM, which was then further diluted with 1.5 mL OptiMEM / PEI. DNA / PEI mixtures were incubated for 15 minutes at 25$^{o}$C prior to addition to cell cultures (100 uL / well in biological triplicates = 1 µg DNA / well; 3 mL / 10 cm dish = 10 µg DNA / 10 cm dish). Sixteen hours post-transfection, DMEM growth medium lacking geneticin was replaced and cells were cultured for 72 hours until experimental or lentivirus harvesting. ACH2 and U1 cells were seeded in 90 µL at 5 x $10^4$ cells / well in round-bottom 96-well culture plates 24 hours prior to transfection with pLentiCRISPR-RFP657. pLenti-CRISPR-RFP657 plasmid (1.08 µg ) was diluted in 81 uL OptiMEM, then supplemented with 3.24 µL TransIT®-Jurkat transfection reagent. After 20 minutes incubation at 25$^{o}$C, 10 uL of the transfection mixtures were added per well in biological replicates (n = 8) and cultured for 72 hours followed by chemical stimulation to assay viral outgrowth.

**Lentivirus Production and Transduction**

Supernatant from pHIV-1 transfected HEK293FT cells was collected 72 hours post-media exchange, centrifuged at 650 RCF for 10 minutes to remove cellular debris and filtered through 0.45 µm PVDF membranes to exclude large exosomes and proteins. HIV-1 was concentrated by layering 10 mL of the filtered supernatant on 2.5 mL 10% w/v sucrose in 50 mM Tris-HCl, 100 mM NaCl, 0.5 mM EDTA, followed by ultracentrifugation at 112,000 RCF for 3.5 hours at 4$^{o}$C. Resulting pellets were resuspended in 250 uL phosphate buffered saline (PBS), aliquoted at 50 uL per tube, and stored at -80$^{o}$C. HIV-1 viral stocks were titered using the Lenti-X™ qRT-PCR Titration Kit according to the manufacturer's protocol. Preparation of third generation CRISPR-transducing lentivirus using our cloned pLentiCRISPR-RFP657 was commercially outsourced to the University of Iowa Viral Vector Core Facility (https://medicine.uiowa.edu/vectorcore/) and titered using digital droplet PCR. U1 or ACH2 cells were seeded at 3 x $10^4$ cells / well in round-bottom 96-well culture plates 24 hours prior to transduction, in the absence or presence of IL-7 (5 ng / mL) mitogen, respectively. Virus was diluted in serum-free RPMI containing 8 µg / mL polybrene. Multiplicities of infection (MOIs) were calculated as viral RNA copies / cell for HIV-1 infection and transducing units / cell for CRISPR-transduction. Wells were infected or transduced using 50 µL diluted virus / well and spin-inoculated at 2400 RCF for 2 hours at 25$^{o}$C. Microtiter plates were then incubated for 4 hours at 37$^{o}$C / 5% $CO_2$ and supplemented with 100 µL growth media. Sixteen hours thereafter, cells were washed 3 times in PBS and cultured in 100 µL growth medium, which was exchanged every 48 hours until experimental termination.

**Viral Outgrowth Assays**

Transfected or transduced latently infected ACH2 and U1 cells were stimulated with 15 ng / mL TNFα (ACH2 cells) or 20 ng / mL phorbol 12-myristate 13-acetate (PMA; U1 cells) diluted in growth media in biological quadruplicates. Biological quadruplicates cultured in growth media alone served as unstimulated controls. Cells were further incubated for 72 hours until terminal harvest.

**Nucleic Acid Isolation**
Biological replicates were pooled and fractioned for DNA and RNA extraction. DNA was extracted using NucleoSpin Tissue XS Micro kit (Machery-Nagel) according to the manufacturer's protocol. TRIzol (Invitrogen) RNA extraction was performed beginning with addition of 600 µL TRIzol to cell pellets followed by mechanical lysis at 30 Hz for 10 minutes using a TissueLyser II instrument (Qiagen). Subsequent steps were performed according to the TRIzol User Guide (https://assets.thermofisher.com/TFS-Assets/LSG/manuals/trizol_reagent.pdf) with volumes scaled proportionately to account for the initial 600 µL TRIzol.

**Polymerase Chain Reaction (PCR)**
CRISPR-mediated excision of proviral DNA was determined by PCR using 150 ng template DNA, 12.5 µL PrimeTime Gene Expression Master Mix (IDT), 1 µL of each 10 µM forward and reverse primers, and volume adjusted to 25 µL with water. Conventional PCR was performed using thermal cycling under the following conditions: 95°C for 3 minutes; 35 cycles of 95°C for 5 seconds, 61.4°C for 30 seconds, 72°C for 30 seconds; and 72°C for 5 minutes. Nested PCR was performed by subjecting DNA from latently infected ACH2 or U1 cells to 15 rounds of PCR as described above. Contents were then diluted 1:10 in water and 2 µL thereof served as template to 30 additional rounds of PCR using nested primers 300b and 301b according to identical thermal cycling conditions. Off-target PCRs were performed using 150 ng template DNA extracted from HEK293FT transfected cells, 12.5 µL AmpliTaq Gold 360 master mix, 2.5 µL GC enhancer, 1 µL of each forward and reverse primers (10 µM), and volume adjusted to 25 µL with water. Thermal cycler conditions were as follows: 95°C for 10 minutes; 35 cycles at 95°C for 30 seconds, 52-60°C (depending on primer Tm) for 30 seconds, and 72°C for 30 seconds; and 72°C for 7 minutes. Sequences of PCR primers are provided in **Table S6**. PCR contents were run in 1% agarose / TAE gels by electrophoresis at 75V for 35 minutes. Gel-extracted PCR amplicons and total PCR contents were Sanger sequenced and subjected to Synthego Inference of CRISPR Edits (ICE) Analysis v2.0 (https://ice.synthego.com/#/; accessed March – November 2020).

**RT-qPCR**
For quantitative reverse transcriptase (RT) PCR (RT-qPCR), 10 ng RNA was first digested with DNase I (Zymo Research). Reverse transcriptase reactions were carried out (42°C for 30 minutes, 95°C for 2 minutes) using Verso cDNA Synthesis Kit (Thermo Scientific) in a volume of 10 µL using 1 µL of DNase I (Zymo Research)-treated RNA and random hexamers. RT-qPCR was subsequently performed to measure *gapdh* or *spCas9* expression using standard curve-based absolute quantitation. A standard curve ($10^5$-1 copy) was generated for *gapdh* using DNA collected from untreated ACH2 cells using a conversion of 1 diploid genome = 6.6 picograms. An *spCas9* standard curve ($10^9$-$10^3$ copies) was made using pLenti-CRISPR-RFP657 using the plasmid molecular weight of $7.22 \times 10^6$ Da to calculate that $10^9$ copies equal 12.01 ng. Ten microliter *spCas9* RT-qPCR reactions were prepared using 1 µL cDNA template, 5 µL PrimeTime Gene Expression Master Mix (IDT), 0.5 µL of each 10 µM forward and reverse primers, and 0.2 µL 10 µM TaqMan probe. RT-qPCR for reference gene were similarly carried out using 0.5 µL Human GAPDH Endogenous Control (Applied Biosystems). All RT-qPCR reactions were performed in technical triplicates under the following thermal cycling conditions: 95°C for 3 minutes, 60 cycles of 95°C for 15 seconds followed by 60°C for 1 minute. Primers and probe sequences are provided in **Table S4**.

**RT Activity Assay**
In 96-well round-bottom plates, 10 µL of culture supernatants from biological replicates (n ≥ 3) were added to 10 µL RT solution A (100 mM Tris-HCl pH 7.9, 300 mM KCl, 10 mM DTT, 0.1% NP-40) then incubated for 15 minutes at 37°C / 5% $CO_2$. Twenty-five microliters of RT solution B (50 mM Tris-HCl pH 7.9, 150 mM KCl, 5 mM DTT, 15 mM $MgCl_2$, 0.05% NP-40, 0.250 U/mL oligo dt pd(T)$_{12\text{-}18}$) freshly supplemented with 10 µCi / mL $^3$H-TTP (4 µL / mL; $^3$H-deoxythimidine 5'-triphosphate, tetrasodium salt, (methyl-$^3$H)) were then added to each reaction well and incubated 20 hours at 37°C and 5% $CO_2$. Fifteen minutes prior to harvesting, 50 µL of ice cold 10% trichloroacetic acid (TCA) was added. Contents were then vacuum filtered across 96-well MicroHarvest Plates (Perkin Elmer). Plates were subsequently washed 3x with each 10% TCA and 100% ethanol and dried for

5 minutes at 67°C. Twenty-five microliters of Microscint-20 (Perkin Elmer) were pipetted to each well. Plates were read via TopCount Scintillation counter (Perkin Elmer). Radioactive counts per minute (CPM), proportionate to the amount of virus contained in sample culture supernatant fluids, were normalized to trypan blue-based cell counts taken at the time of harvest.

**Flow Cytometry**

CEMss CD4+ T cells infected with GFP-reporting HIV-1$_{NL4\text{-}3\text{-}\Delta tat\text{-}\Delta nef\text{-}eGFP}$ were washed once with PBS and fixed in 2% paraformaldehyde (PFA). GFP expression was measured in fixed cells using an Attune NXT Flow Cytometer (Thermo Fisher) and reported as % of gated single-cell population.

**Statistical Analyses**

Experiments were run with biological replicates (n = 3 or 4) in at least two separate trials, as indicated in figure legends. RT activity assays and RT-qPCR were further performed with technical triplicates. Data are presented as mean ± standard error of the mean (SEM). Pearson correlation between mean RT activity reductions and percent gRNA conservation was performed using two-tailed p-value determination. Statistical differences between unstimulated vs stimulated RT activity values and stimulated control vs stimulated treatment RT activity values were calculated with two-way ANOVA by Sidak correction for multiple comparisons. One-way ANOVA with Dunnett correction for multiple comparisons was employed to assess for differences between the stimulation indices of control- and *tat*-targeted treatments. All statistical calculations set type I error cutoff $\alpha = 0.05$ and were performed using GraphPad Prism v7.0a for Mac OS X (GraphPad Software, San Diego, California USA, www.graphpad.com).

**Data Availability**

DNA Sanger sequence chromatograms and spreadsheets used for Inference of CRISPR Edits v2 (ICEv2), along with algorithm results, are uploaded to Mendeley Data repository are accessible at doi:10.17632/phyy89w9c2.1 (embargoed until publication); https://data.mendeley.com/datasets/phyy89w9c2/draft?a=2d79cf40-8783-457d-86f1-8c1662877a8a

**Supplementary Figures.**

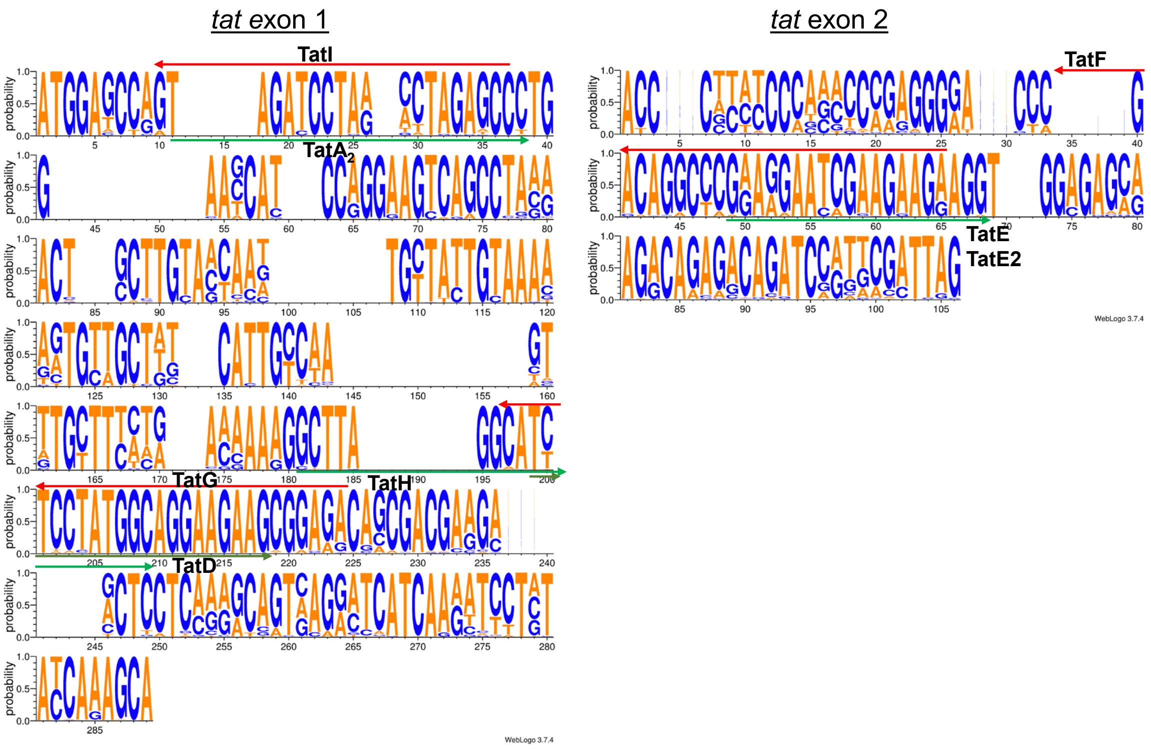


**Fig. S1.** CRISPR-Cas9 Targeting HIV-1 *tat* Consensus Sequence. Sequence logos depicting loci of spCas9 gRNA targeting HIV-1 *tat* exon 1 (nucleotide positions 5831-6045 using HIV-1HXB2 reference strain) and *tat* exon 2 (nucleotide positions 8379-8469) amongst all curated HIV-1 strain sequences compiled as of 2018 (n = 4004; http://www.hiv.lanl.gov/) were generated using WebLogo v3.7.4. Conserved nucleotides are represented as taller letters whereas blank spaces denote frequent gaps in alignment. gRNA directed against sense or antisense strands are shown as rightward (green) or leftward (red) facing arrows, respectively.

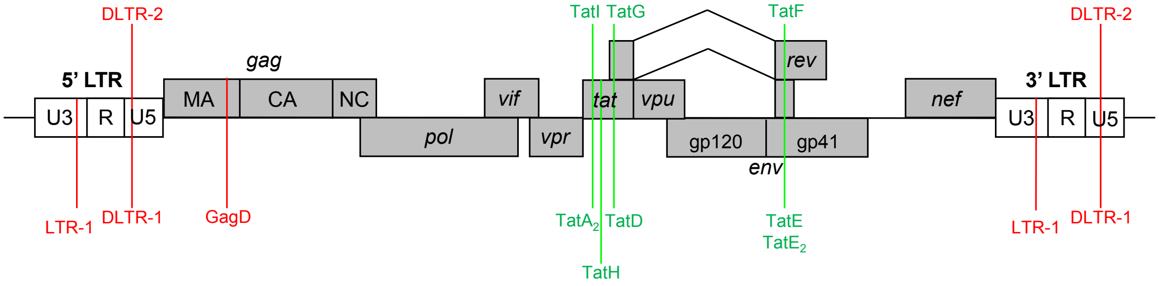


**Fig. S2.** Loci of Experimental spCas9 gRNAs. Guide RNAs directed against HIV-1 LTR / *gag* (red) or *tat* exons (green) overlapping with up to two additional genes were designed using and HIV-1 consensus sequence. The gRNA library was individually or dually cloned into an spCas9 expression vector.

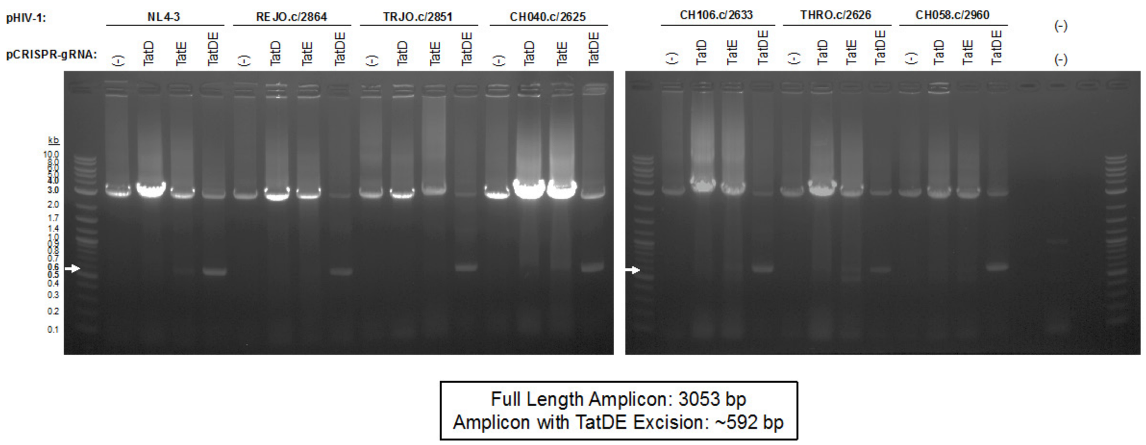


**Fig. S3.** TatDE Dual gRNA Facilitate Multistrain HIV-1 Excision. HEK293FT cells co-transfected with pHIV-1 in the absence or presence of CRISPR-gRNA containing plasmids were assayed by PCR for DNA excision of 2.461 kb between intervening protospacers contained in *tat/rev*. Arrow indicates expected molecular size of TatDE excision band.

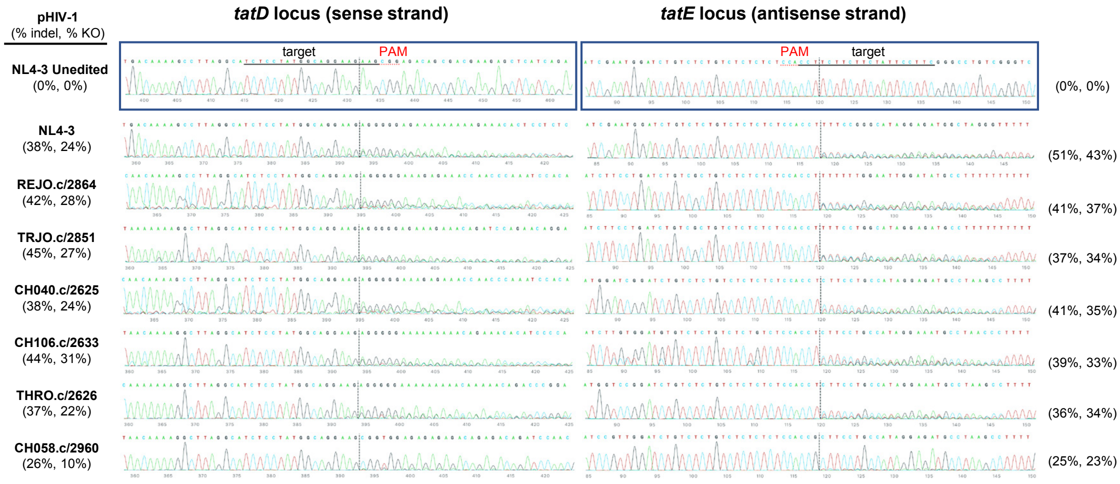


**Fig. S4.** CRISPR Editing at TatD and TatE Loci in Co-Transfection Screen. PCR amplicons of CRISPR and HIV-1 encoding plasmid co-transfections of HEK293FT cells were gel extracted and Sanger sequenced. Chromatograms are accompanied by indel and knockout (KO) percentages as determined by Synthego Inference of CRISPR Edits v2.0 (ICE) algorithm.

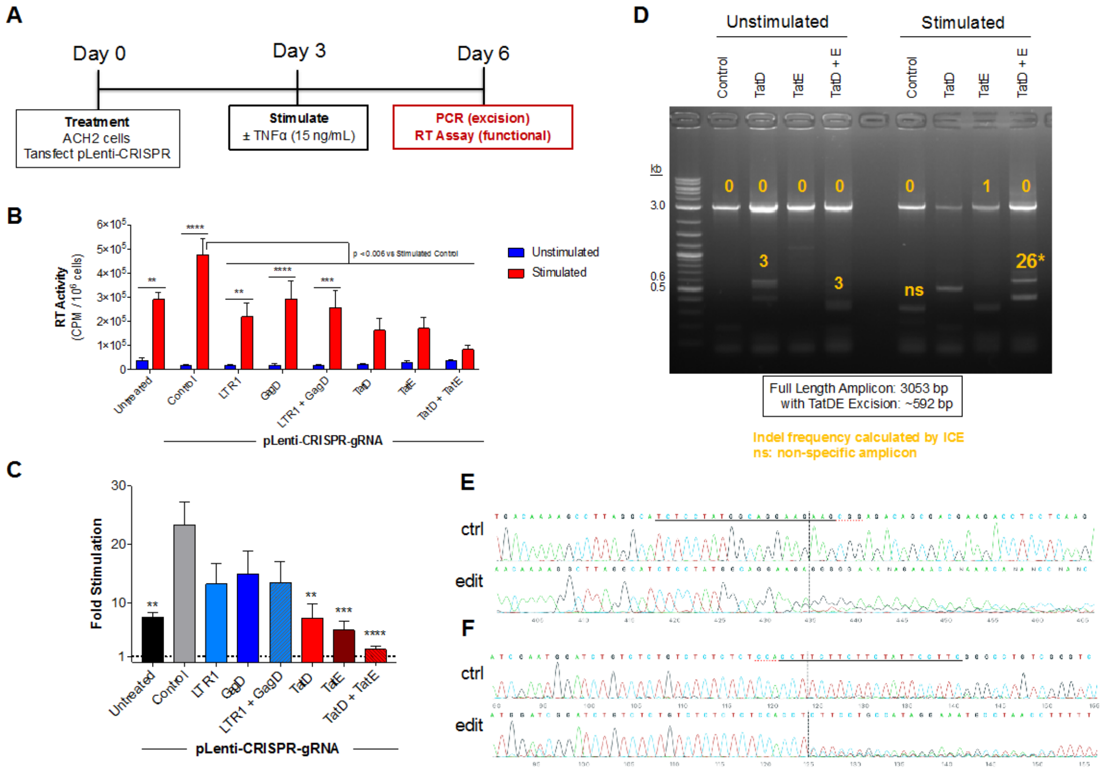


**Fig. S5.** TatDE CRISPR Co-transfection Abolishes Latent HIV-1 in Infected ACH2 T cells. *(A)* ACH2 T cells containing 1 copy of integrated HIV-1 proviral DNA were transfected with 120 ng of CRISPR-Cas9 encoding lentiviral plasmids then stimulated and harvested according to the diagram above. *(B)* Supernatants were measured by RT activity assay 72 hours post stimulation. *(C)* The fold stimulation in *(B)* was determined as the ratio of RT activity in the presence:absence of TNFα. *(D)* DNA was PCR amplified to survey for shortened amplicons resulting from CRISPR-Cas9 mediated excision of intervening DNA. Bands were gel extracted and subjected to ICE to determine indel frequency. *(E-F)* Excision bands from TatD + TatE co-transfection in the presence of TNFα were Sanger sequenced to confirm editing in the TatD *(E)* and TatE *(F)* loci. Data in *(B-C)* depict mean ± SEM from three independent experiments each containing biological triplicates. Significance assessed by two- *(B)* or one-way *(C)* ANOVA.

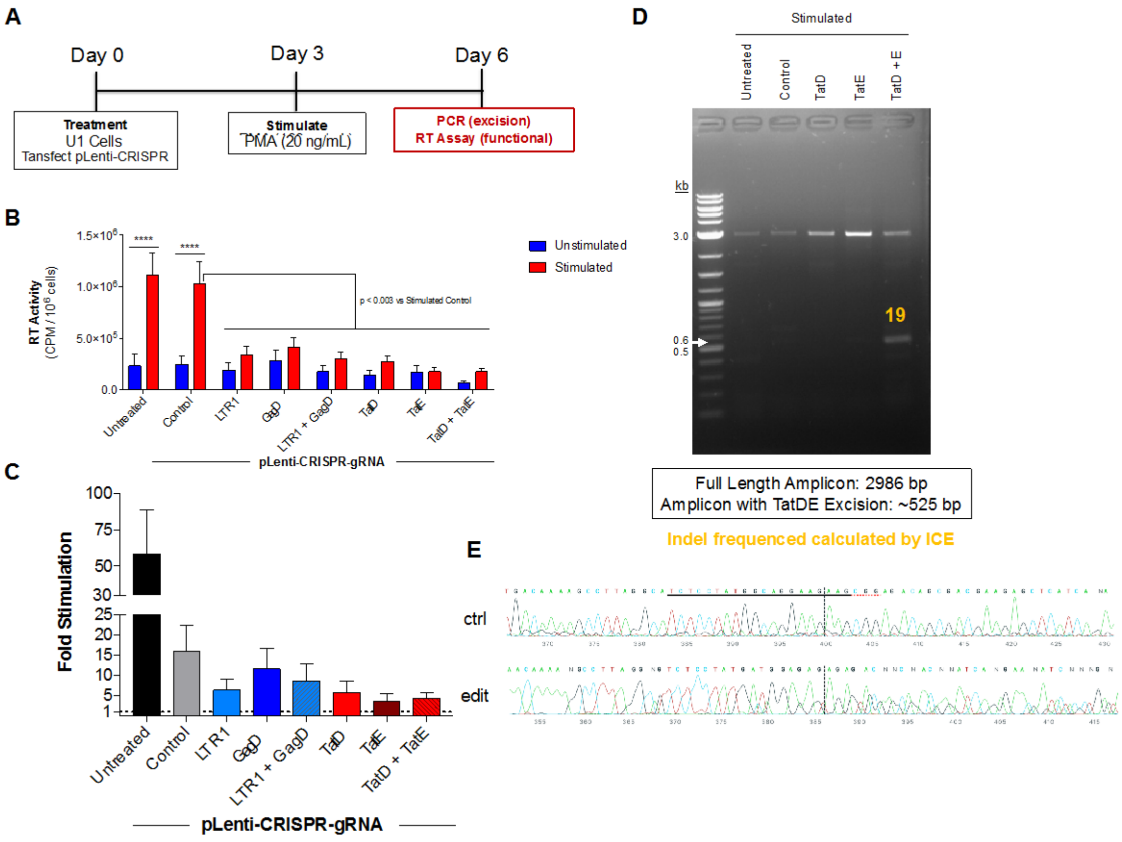


**Fig. S6.** TatDE CRISPR Co-transfection Reduces HIV-1 in Latently Infected U1 Promonocytes. *(A)* U1 promonocytes containing 1-2 copies of integrated HIV-1 proviral DNA were transfected with 120 ng of CRISPR-Cas9 encoding lentiviral plasmids then stimulated and harvested according to the diagram above. *(B)* Supernatants were measured by RT activity assay 72 hours post stimulation. *(C)* The fold stimulation in *(B)* was determined as the ratio of RT activity in the presence:absence of phorbol 12-myristate 13-acetate (PMA). *(D)* Nested PCR was performed to survey for shortened amplicons resulting from CRISPR-Cas9 mediated excision of intervening DNA. Bands were gel extracted and subjected to ICE to determine indel frequency. *(E)* Excision bands from TatD + TatE co-transfection in the presence of PMA were Sanger sequenced to confirm editing in the TatD locus. Data in *(B-C)* depict mean ± SEM from two independent experiments each containing biological triplicates. Significance assessed by two-way ANOVA.

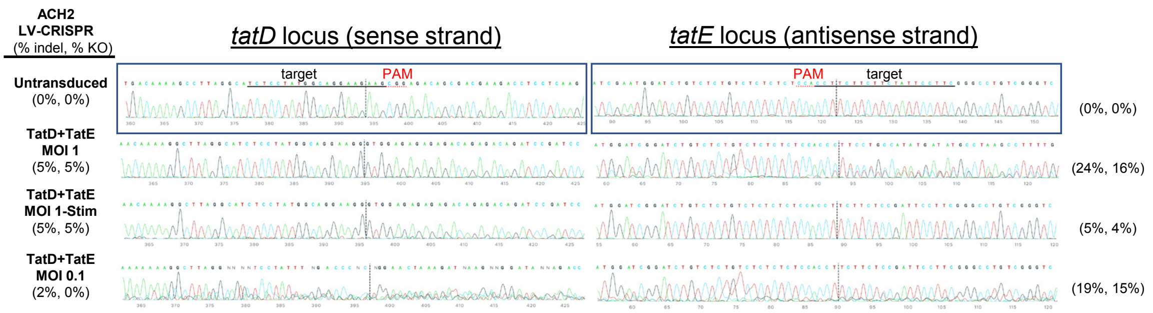


**Fig. S7.** CRISPR Editing at TatD and TatE Loci in Lentiviral-transduced ACH2 T cells. PCR amplicons of ACH2 T cells lentivirally transduced with CRISPR were gel extracted and Sanger sequenced. Chromatograms are accompanied by indel and knockout (KO) percentages as determined by Synthego Inference of CRISPR Edits v2.0 (ICE) algorithm.

**Supplementary Tables.**

**Table S1. Experimental gRNA Targets and Specificities**

| gRNA Name | Target DNA Sequence (5' → 3') | % Sequence Conservation (n = 4004) | CRISPR*spec* Score† |
|---|---|---|---|
| LTR-1 (control) | GCAGAACTACACACCAGGGCC | 27.12 % | 6.937 |
| GagD (control) | GATAGATGTAAAAGACACCA | 11.14 % | 4.071 |
| DLTR-1 (control) | GACTGCTTAAGCCTCAATAA | 29.57 % | 3.364 |
| DLTR-2 (control) | GCTTTATTGAGGCTTAAGCAG | 28.94 % | 2.266 |
| TatG | TCTCCGCTTCTTCCTGCCAT | 67.15 % | 3.803 |
| TatD | TCTCCTATGGCAGGAAGAAG | 59.04 % | 6.284 |
| TatH | GCTTAGGCATCTCCTATGGC | 51.12 % | 4.662 |
| $TatA_2$ | TAGATCCTAACCTAGAGCCC | 29.79% | 7.327 |
| TatI | GGCTCTAGGTTAGGATCTAC | 25.90 % | 6.844 |
| TatE | GAAGGAATCGAAGAAGAAGG | 15.26 % | 3.090 |
| TatF | CCGATTCCTTCGGGCCTGTC | 6.19 % | 6.467 |
| $TatE_2$ | GAAAGAATCGAAGAAGGAGG | 5.82 % | 6.021 |

†CRISPR*spec* score represents the specificity of the selected gRNA for HIV-1 based on a predefined off-target landscape within the human genome. Calculated using CRISPRoff webserver v1.1.

**Table S2. Mosaic gRNA Coverage**

| gRNA Name | % Sequence Conservation (n = 4004 HIV-1 Strains) | Excised Base Pairs | Exons Disrupted |
|---|---|---|---|
| TatD | 59.04 % | - | 2 |
| TatH§ | 51.12 % | - | 1 |
| TatE | 15.26 % | - | 3 |
| TatDH‡ | 59.44 % | 29 | 3 |
| TatDE | 62.19 % | 2442 | 5 |
| TatEH | 56.24 % | 2451 | 4 |

§ 4 base pairs of TatH gRNA overlap with *rev*. Given that CRISPR cleavage, as determined by ICE, occurs 5+ bp distal to PAM, TatH was considered to disrupt 1 exon ($tat_1$).

‡ Although TatDH is a composite of 2 highly conserved gRNAs, they overlap in target recognition and therefore multiplexing did not significantly increase breadth of coverage.

**Table S3. Off-Target TatDE CRISPR Editing**

| gRNA | Guide Target (Protospacer mismatches) + PAM | *Gene* (Locus) | Treatment | % Indel (per pHIV-1 Strain) pCH040.c /2625 | pCH106.c /2633 | pTHRO.c /2626 | pCH058.c /2960 | Gene Function |
|---|---|---|---|---|---|---|---|---|
| TatD | CCTCCCATGGCAGAAAGAAGTGG | *ALDH3B1* (11q13.2) | TatD | 0% | 0% | 0% | 0% | Aldehyde dehydrogenase aids in alcohol metabolism and lipid peroxidation (Accession NG_012282) |
| | | | TatDE | 0% | 0% | 0% | 0% | |
| | CTTCCCACGGCAGGAAGAACCGG | *GPC6* (13q31.3-q32.1) | TatD | 0% | 0% | 0% | 0% | Glypicans encode cell surface coreceptors for control of cell growth and division (Accession NG_011880) |
| | | | TatDE | 0% | 0% | 0% | 0% | |
| | CTTCCTAGGGCAGGAAGAAGGGG | *IGFBP5* (2q35) | TatD | 0% | 0% | 0% | 0% | Insulin-like growth factor-binding protein 5; function in humans under characterized (Accession AC007563) |
| | | | TatDE | 0% | 0% | 0% | 0% | |
| | TTCTCCTAGGAAGGAGGAAGAGG | Intronic (12q24.33) | TatD | 0% | 0% | 0% | 0% | (Accession AC127071) |
| | | | TatDE | 0% | 0% | 0% | 0% | |
| | AATCTTATGGCAGGAAGAAGAGG | Intronic (9q22.2-q31.1) | TatD | 0% | 0% | 0% | 0% | (Accession AL137847) |
| | | | TatDE | 0% | 0% | 0% | 0% | |
| TatE | GAAGAAAACAAAGAAGAAGGAGG | Intronic (18q12.2) | TatE | 0% | 0% | 0% | 0% | (Accession AC118757) |
| | | | TatDE | 0% | 0% | 0% | 0% | |
| | GAAGGAAGAGAAGAAGAAGGAGG | Intronic (19q13.33) | TatD | 0% | 0% | 0% | 0% | (Accession AC010330) |
| | | | TatDE | 0% | 0% | 0% | 0% | |
| | GAAGGAATCA-AGAAGAAGGAGG | Intronic (4q35.1) | TatD | 0% | 0% | 0% | 0% | (Accession AC079080) |
| | | | TatDE | 0% | 0% | 0% | 0% | |
| | GAAAGGAAGGAAGAAGAAGGGGG | Senescence gene region (8p11.2) | TatD | 0% | 0% | 0% | 0% | (Accession AP000078.1) |
| | | | TatDE | 0% | 0% | 0% | 0% | |
| | AAAGGAATCGAAGAAGAAGGGGG | Intronic (7q31.33) | TatD | 0% | 0% | 0% | 0% | (Accession AC005521) |
| | | | TatDE | 0% | 0% | 0% | 0% | (Accession AC005521) |

**Table S4: Off-Target LTR1 / GagD (LG) CRISPR Editing**

| gRNA | Guide Target (Protospacer mismatches) + PAM | *Gene* (Locus) | Treatment | % Indel (per pHIV-1 Strain) pCH040.c /2625 | pCH106.c /2633 | pTHRO.c /2626 | pCH058.c /2960 | Gene Function |
|---|---|---|---|---|---|---|---|---|
| LTR-1 | GCAGAACCACTCCCCAGGGCCTGG | *GRIK3* (1p35.1) | LTR-1 | 0% | 0% | 0% | 0% | Glutamate ionotropic receptor kainate type subunit 3; glutamate neurotransmitter receptor subunit. (NG_011447.1) |
| | | | LG | 0% | 0% | 0% | 0% | |
| | AAAGAACAACACACCAGGGAGGGG | *CADPS2* (7q31.31) | LTR-1 | 0% | 0% | 0% | 0% | Calcium dependent secretion activator 2; regulates exocytosis of synaptic vesicles in neurons and neuroendocrine cells. (NG_016215) |
| | | | LG | 0% | 0% | 0% | 0% | |
| | TAAAAACTACACACAAGGGCCAGG | Intronic (3q13.32) | LTR-1 | 0% | 0% | 0% | 0% | (Accession AC021889) |
| | | | LG | 0% | 0% | 0% | 0% | |
| | AGGGAAACACACACCAGGGCCAGG | *GP6C* (13q31.1) | LTR-1 | 0% | 0% | 0% | 0% | Glypican 6; glycosphlphophatidylinositol-anchored proteoglycan implicated in cell growth and cell division. (NG_011880.1) |
| | | | LG | 0% | 0% | 0% | 0% | |
| | TAGAACTACACCACCAGGGCCCGG | *CRX* (19q13.33) | LTR-1 | 0% | 0% | 0% | 0% | Cone-rod homeobox; plays role in differentiation of photoreceptor cells and maintenance of cone & rod function. (AC008745.7) |
| | | | LG | 0% | 0% | 0% | 0% | |

| | | | | | | | | |
|---|---|---|---|---|---|---|---|---|
| **GagD** | CACAGAGGCAAAAGACACCATGG | otopetrin 1 pseudogene (2p11.2) | GagD | 0% | 0% | 0% | 0% | Nonfunctional DNA segment derived from *OTOP1* (4p16.3), a gene that is required for the formation of inner ear otoliths. (NG_021568.4) |
| | | | LG | 0% | 0% | 0% | 0% | |
| | GATAGATGTAAAAGACACTGTGG | *ZNF420* (19q13.41) | GagD | 0% | 0% | 0% | **n.d.** | Zinc finger protein 420; negatively regulates p53-mediated apoptosis especially in testes and ovaries. (AC008733.9) |
| | | | LG | 0% | 0% | 0% | **n.d.** | |
| | GATAGATGTAAAAGACACCAGGG | *MLLT3* (9p22.1) | GagD | 0% | 0% | 0% | 0% | MLLT3 super elongation complex unit. (AK512635.8) |
| | | | LG | 0% | 0% | 0% | 0% | |
| | GAGAAATATAAAAGACACCATGG | Intronic (5q22.2) | GagD | 0% | **n.d.** | 0% | 0% | (Accession AC137549.1) |
| | | | LG | 0% | **n.d.** | 0% | 0% | |
| | TATAGCTGTAAAAACACCA-AGG | *PTPRG* (3p13.2) | GagD | 0% | 0% | 0% | 0% | Protein tyrosine phosphatase receptor type G; signaling molecule in mitosis, differentiation, and oncogenic transformation. (AC092502.2) |
| | | | LG | **n.d.** | 0% | 0% | 0% | |

**n.d**. no data available

**Table S5. Cloning Primers**

| Vector | gRNA | Forward Oligo (5'→3')[a] | Reverse Oligo (5'→3')[b] |
|---|---|---|---|
| px333 | LTR-1 | caccCAGAACTACACACCAGGGCC | aaacGGCCCTGGTGTGTAGTTCTG |
| | GagD | caccGATAGATGTAAAAGACACCA | aaacTGGTGTCTTTTACATCTATC |
| | DLTR-1 | caccGACTGCTTAAGCCTCAATAA | aaacTTATTGAGGCTTAAGCAGTC |
| | DLTR-2 | caccGCTTTATTGAGGCTTAAGCAG | aaacCTGCTTAAGCCTCAATAAAGC |
| | TatA2 | caccTAGATCCTAACCTAGAGCCC | aaacGGGCTCTAGGTTAGGATCTA |
| | TatD | caccTCTCCTATGGCAGGAAGAAG | aaacCTTCTTCCTGCCATAGGAGA |
| | TatE | caccGAAGGAATCGAAGAAGAAGG | aaacCCTTCTTCTTCGATTCCTTC |
| | TatE2 | caccGAAAGAATCGAAGAAGGAGG | aaacCCTCCTTCTTCGATTCTTTC |
| | TatF | caccCCGATTCCTTCGGGCCTGTC | aaacGACAGGCCCGAAGGAATCGG |
| | TatG | caccTCTCCGCTTCTTCCTGCCAT | aaacATGGCAGGAAGAAGCGGAGA |
| | TatH | caccGCTTAGGCATCTCCTATGGC | aaacGCCATAGGAGATGCCTAAGC |
| | TatI | caccCCGATTCCTTCGGGCCTGTC | aaacGACAGGCCCGAAGGAATCGG |
| pLentiCRISPR-RFP657 | Control | caccGGAGACGTTTGTACGTCTCT | aaacAGAGACGTACAAACGTCTCC |
| | TatD | caccGTCTCCTATGGCAGGAAGAAG | aaacCTTCTTCCTGCCATAGGAGAC |
| | TatE | caccGGAAGGAATCGAAGAAGAAGG | aaacCCTTCTTCTTCGATTCCTTCC |
| pHIV-1NL4-3-Δnef-eGFP | ΔTatD | GAAGCGGGGGAGACAGCGACGAAGAGCTC | TGTCTCCCCCGCTTCTTCCTGCCATAG |
| | ΔTatE | TAGAAGAAAGAGACAGAGACAGATCCATTCG | TGTCTCTTTCTTCTATTCCTTCGGGCC |

[a] Overhang sequences complementary to digested vector represented as lower case “cacc”
[b] Overhang sequences complementary to digested vector represented as lower case “aaac”

**Table S6: PCR Primers**

| | Forward Oligo | | | Reverse Oligo | | |
|---|---|---|---|---|---|---|
| **PCR** | **Template DNA (locus)** | **Primer Name** | **Sequence (5'→3')** | **Template DNA (locus)** | **Primer Name** | **Sequence (5'→3')** |
| TatDE Excision | HIV-1$_{HXB2}$ (5518-5537)<br>HIV-1$_{CH040.c/2625}$ (5514-5533)<br>HIV-1$_{CH106.c/2633}$ (5515-5534)<br>HIV-1$_{TRJO.c/2851}$ (5542-5561)<br>HIV-1$_{CH058.c/2960}$ (5535-5554) | Primer 300 | AAGCCACCTTTGCCTAGTGT | HIV-1$_{HXB2}$ (8560-8580)<br>HIV-1$_{CH040.c/2625}$ (8525-8545)<br>HIV-1$_{THRO.c/2626}$ (8643-8663)<br>HIV-1$_{REJO.c/2864}$ (8576-8596)<br>HIV-1$_{TRJO.c/2851}$ (8586-8606) | Primer 301 | CCAGAAGTTCCACAATCCTCG |
| | HIV-1$_{THRO.c/2626}$ (5549-5568) | Primer 302 | AAGCCGCCTTTGCCTAGTGT | HIV-1$_{CH058.c/2960}$ (8546-8566) | Primer 303 | CCAGAAGTTCCACAGTCCTCG |
| | HIV-1$_{REJO.c/2864}$ (5538-5557) | Primer 304 | AAGCCACCTTTGCCTAGTAT | HIV-1$_{CH106.c/2633}$ (8580-8600) | Primer 305 | CCAGAAGTTCCACTATCCTCG |
| TatDE Excision (nested) | HIV-1$_{HXB2}$ (5557-5578)<br>HIV-1$_{CH040.c/2625}$ (5553-5574)<br>HIV-1$_{CH106.c/2633}$ (5554-5575)<br>HIV-1$_{THRO.c/2626}$ (5588-5609)<br>HIV-1$_{REJO.c/2864}$ (5577-5598)<br>HIV-1$_{TRJO.c/2851}$ (5581-5602)<br>HIV-1$_{CH058.c/2960}$ (5574-5595) | Primer 300b | AGATGGAACAAGCCCCAGAAGA | HIV-1$_{HXB2}$ (8526-8552)<br>HIV-1$_{CH040.c/2625}$ (8491-8517)<br>HIV-1$_{CH106.c/2633}$ (8546-8572)<br>HIV-1$_{THRO.c/2626}$ (8609-8635)<br>HIV-1$_{REJO.c/2864}$ (8542-8568)<br>HIV-1$_{TRJO.c/2851}$ (8552-8578)<br>HIV-1$_{CH058.c/2960}$ (8512-8538) | Primer 301b | CAAGAGTAAGTCTCTCAAGCGGTGGTA |
| TatD Off-Target | chr11: 68022307-68022329 | Primer 182-D2 Off F | TGAACTCCAAGGGTTCTAAGAT | chr11: 68022733-68022751 | Primer 183-D2 Off R | TTGGAGAAGGCGTACAGG |
| | chr13: 93771074-93771102 | Primer 194-D3 Off F | GTGCCAAACATATTGATAACACATACAC | chr13: 93771551-93771574 | Primer 195-D3 Off R | CTCCTCAAAGCTCACCAAATTCC |

| | | | | | | |
|---|---|---|---|---|---|---|
| | chr2: 216741039-216741061 | Primer 274-D7 Off F | GTTTATAATTGCAGAAGCACTG | chr2: 216741522-216741539 | Primer 275-D7 Off R | AACCCAATTCTGTCCGA |
| | chr12: 130325120-130325144 | Primer 184-D9 Off F | AAAGTGGATGACAAGTTCTATCAG | chr12: 130325598-130325620 | Primer 185-D9 Off R | AGACTTGAGATGAGAGAAACCC |
| | chr9: 83240840-83240862 | Primer 247-D10 Off F | TCAAGGTTCTAAGTTTCTGGCA | chr9: 83241314-83241340 | Primer 248-D10 Off R | CCAGAACATAAGCTGTAATGTAAACC |
| TatE Off-Target | chr18: 35964872-35964889 | Primer 282-E3 Off F | TTCAGATGCTGGAGTAG | chr18: 35965355-35965372 | Primer 283-E3 Off R | TTAGCTGGACTTGATGG |
| | chr19: 47991242-47991260 | Primer 278-E4 Off F | GACCCAAACCAGCAGGAG | chr19: 47991721-47991742 | Primer 279-E4 Off R | GGAGGAGGTGAAGAAGGAAGA |
| | chr4: 184272401-184272421 | Primer 226-E7 Off F | CACACAGAGGCTTTGGAACT | chr4: 184272881-184272901 | Primer 227-E7 Off R | ACTATGTACCCAGGCAGCTA |
| | chr8: 36453259-36453280 | Primer 244-E12 Off F | GTACCTTAGAGGTTTCCAATC | chr8: 36453739-36453759 | Primer 245-E12 Off R | TGAAGAATCAAATGCAACAC |
| | chr7: 124503052-124503072 | Primer 373-E14 Off F | CCACAGTTTATATATTCCAC | chr7: 124503057-124503075 | Primer 374-E14 Off R | CTTGAAGCCACCTTATAC |
| LTR-1 Off-Target | chr1: 36971727-36971747 | Primer 254-LTR1 Off F | AAGACTTCTAGGTGGATAGC | chr1: 36972189-36972207 | Primer 255-LTR1 Off R | CCTGTGGATTCTCCTTCA |
| | chr7: 122425478-122425495 | Primer 256-LTR2 Off F | GTGGTCCCAAGCTACTC | chr7: 122425924-122425943 | Primer 257-LTR2 Off R | GAGTTAAAGCAAGGGTCAG |
| | chr3: 117193688-117193710 | Primer 258-LTR3 Off F | GTATTGCTTTAGACTCCTAAGT | chr3: 117194162-117194179 | Primer 259-LTR3 Off R | CAAGATCTCGGCTCACT |
| | chr13: 94251055-94251072 | Primer 260-LTR4 Off F | CCCAAATGCCCATCAAT | chr13: 94251526-94251545 | Primer 261-LTR4 Off R | CACAAGTTCTGGGATTTGG |
| | chr19: 47786499-47786517 | Primer 262-LTR11 Off F | TTACAGGCATATGTCACC | chr19: 47786908-47786928 | Primer 263-LTR11 Off R | CTTGCTTAAACAAAGAACTG |
| GagD Off-Target | chr2: 91614689-91614709 | Primer 264-Gag1 Off F | GGCATCACATCTGGGTTCAA | chr2: 91615200-91615221 | Primer 265-Gag1 Off R | CTTTCTGCTTCTGGCCCTATC |
| | chr19: 36832404-36832427 | Primer 367-Gag12 Off F | AGAATAGTAGTGATGCTGAACA | chr19: 36832409-36832427 | Primer 368-Gag12 Off R | CCAGGGCTGCTATCATTT |
| | chr9: 20223152-20223174 | Primer 369-Gag13 Off F | AGCAGCATTAAACTTGGAAAGG | chr9: 20223152-20223175 | Primer 370-Gag13 Off R | AGTGTTGAGATTAGAGGTGTGAG |
| | chr5: 112567658-112567678 | Primer 270-Gag4 Off F | TGCTAGGGCAGAGAGGATAA | chr5: 112568112-112568134 | Primer 271-Gag4 Off R | GACGTTCTCATGGAATCTCACA |
| | chr3: 62330371-62330388 | Primer 272-Gag11 Off F | ATATGGCCTGTGACCAA | chr3: 62330762-62330783 | Primer 272-Gag11 Off R | GCTCAGTAAGATACCTCAGTA |
